# Correlation of Plant Bioelectrical Signals with Potential Ionic Energy Flow under Different Stress

**DOI:** 10.64898/2026.08.28.747893

**Authors:** Shilpa Chandra, Chayan Kanti Nandi, Laxmidhar Behera

**Affiliations:** Centre for Indian Knowledge System and Mental Health Applications, Indian Institute of Technology Mandi, HP-175005; School of Chemical Sciences, Indian Institute of Technology Mandi, HP-175005; School of Computing and Electrical Engineering, Indian Institute of Technology Mandi, HP-175005

**Keywords:** Plant electrophysiology, Variation potential, Action potential, Systemic potential, multidomain signal analysis, Ionic flow

## Abstract

All living organisms rely on the movement of ions across cell membranes as the fundamental physical basis of their internal energy and signaling, and plants are no exception. Plants perceive, integrate, and respond to environmental stimuli through electrical signals, classified as action, variation, and system potentials, that are coupled with calcium waves, reactive oxygen species, and hydraulic and hormonal changes to coordinate whole-organism responses despite the absence of a nervous system. Yet most studies characterize these signals using a single feature, such as amplitude or spike duration, in a single tissue, an approach that cannot establish how such signals correspond to the underlying ionic activity, mobility, and structural complexity of the signaling environment, or how this correspondence varies across organs. Here, we correlate plant bioelectrical signals with potential ionic energy flow using a multi-domain framework, combining discrete spike events, continuous waveform properties, spectral composition, and signal complexity applied to leaf, stem, and root recordings from tomato (*Solanum lycopersicum*) exposed to different stimulus. Electrical activity with increased stimulus strength, likely reflecting increased ionic flow, with the root showing the largest response. This suggests plant electrical signaling works as a distributed, ion-based information system, useful for stress monitoring and bio-inspired sensor design.

## Introduction

All living organisms rely on the movement of ions across cell membranes as the fundamental physical basis of their internal energy and signaling, and plants are no exception.^1^ Despite lacking a centralized nervous system, plants perceive, integrate, and respond to environmental stimuli through electrical signals generated by these ionic fluxes, using this ion-based communication as a rapid, whole-body alternative to neuronal signaling. Plant electrical events are commonly categorized into action potentials, variation potentials, and system potentials, each associated with distinct physiological triggers such as mechanical perturbation, wounding, and environmental stress, and each reflecting a characteristic pattern of ion channel activation and membrane depolarization.^2–5^ These signals propagate across tissues and are coupled with calcium waves,^6,7^ reactive oxygen species,^8,9^ hydraulic changes,^10^ and hormonal signaling,^11^ forming a multi-layered ionic and biochemical network that allows local stimuli to generate coordinated responses at the whole-plant level. Together, these ion-mediated processes constitute the physical substrate through which a stimulus detected at one point in the plant is translated into a change in electrical and physiological state elsewhere.

A key distinction within this signaling network lies between short-distance local and long-distance systemic responses. Local signaling involves transient, low-amplitude electrical and ionic changes confined to the site of stimulation, typically triggered by mild stimuli such as touch or light, dissipating rapidly without engaging distant tissue, as seen in the touch-induced leaf-folding response of *Mimosa pudica*^12^ or the rapid trap closure of *Dionaea muscipula* following mechanical stimulation of trigger hairs.^13^ Long-distance signaling, by contrast, involves propagation across multiple organs, mediated by variation and system potentials, triggered by stronger perturbations such as mechanical injury, herbivory, or thermal stress, and coupled with systemic calcium and reactive oxygen species waves that reinforce the electrical signal as it travels,^5^ as in the wound-induced systemic activation of proteinase inhibitor genes in distal, undamaged leaves of tomato following localized insect feeding,^14– 16^ or the glutamate-receptor-like-channel-mediated leaf-to-leaf electrical and calcium signaling triggered by wounding and herbivory.^17,18^ Real-time intracellular recordings using living aphids as bioelectrodes have further shown that such depolarization waves reach neighboring, undamaged tissue within seconds of localized feeding damage.^19^ Plant signaling does not operate as a binary switch between these two regimes but along a continuum, in which increasing stimulus intensity produces progressively broader spatial engagement of the underlying ionic and electrical machinery, from a localized event to whole-plant coordination **(Figure 1)**.^6^ The root apex has specifically been proposed as a site where convergent electrical, chemical, and mechanical signals, and the ionic fluxes underlying them, are coordinated into whole-organism responses, functioning as a possible convergence point for information arriving from multiple tissues, a hypothesis that has not been directly tested using multi-domain electrophysiological data.^20,21^

**Figure 1:**
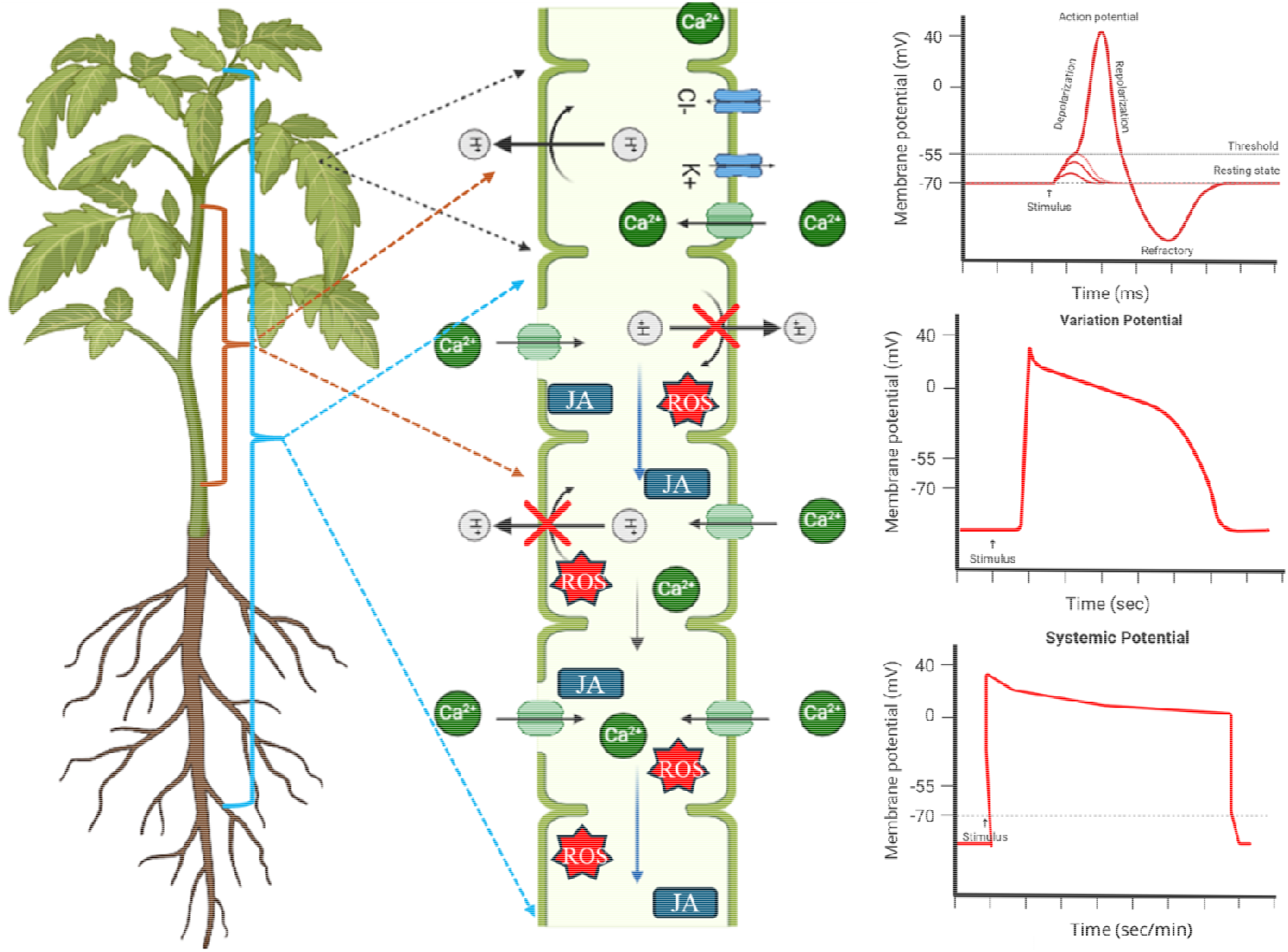
Stimulus-dependent electrical signal propagation in *Solanum lycopersicum*. (Left) Schematic illustrating three long-distance signaling modalities across leaf, stem, and root: action potentials (black arrows), variation potentials (brown dashed arrows), and system potentials (blue dashed arrows), coupled with sequential Ca^2^□ influx, ROS generation, and jasmonic acid (JA) biosynthesis along the vascular axis. (Right) Representative membrane potential waveforms for each signal type: action potential showing rapid depolarization and repolarization on a millisecond timescale; variation potential displaying slow, sustained depolarization on a second timescale driven by hydraulic pressure-coupled ion channel activation; and system potential showing prolonged plateau depolarization reflecting whole-plant apoplastic electrical coordination.

Despite this progress, a central problem remains unresolved. Most electrophysiological studies examine a single stimulus in a single tissue, typically the leaf, and rely on a single feature, such as amplitude or spike duration, to characterize the response, an approach that cannot establish how such signals correspond to the underlying ionic activity, mobility, and structural complexity of the signaling environment, or how this correspondence differs across organs.^22,23^ Yet plant electrical signals plausibly reflect ionic energy flow across several complementary dimensions at once, including discrete event structure, sustained signal energy, spectral composition, and higher-order temporal complexity, none of which is captured by any single metric in isolation, and each of which may index a different aspect of the underlying ionic process.^7^ The novelty of the present work lies in combining these complementary dimensions within a single, organ-resolved framework, allowing the ionic basis of plant electrical signaling to be examined as a coherent, multi-domain system rather than through isolated, single-feature measurements that capture only one facet of the response at a time.

This study systematically recorded electrophysiological responses to six stimuli spanning a wide range of biological significance, touch, sound, smell, red light, cut, and burn, from leaf, stem, and root of tomato (*Solanum lycopersicum*), and correlated each response with potential ionic energy flow using discrete spike events, continuous time-domain waveform properties, frequency-domain spectral organization, and nonlinear Hjorth descriptors of signal complexity. Across all four domains, electrical activity, and the ionic energy flow it reflects, followed a consistent, graded hierarchy, rising from minimal, localized change under mild stimuli to widespread, sustained change under severe thermal stress, with the root consistently showing the broadest ionic engagement of any organ across every domain examined. These results provide a quantitative, multi-domain electrophysiological foundation for correlating plant bioelectrical signals with ionic energy flow, with practical implications for real-time plant stress monitoring, precision agriculture, and the design of bio-inspired sensing systems that draw on the same ion-mediated principles of detection and signal propagation observed in this study **(Table S1)**.

## Results and Discussions

Electrophysiological responses of six-week-old tomato (*Solanum lycopersicum*) leaves, stems, and roots were recorded using the Plant SpikerBox system under touch, cut, burn, sound, smell, and red-light stimuli.^24^ The electrical responses were analyzed across four complementary domains: discrete spike events, continuous waveform characteristics, spectral composition, and nonlinear complexity. ^25,26^ These parameters capture different aspects of ion-mediated electrical activity, including the occurrence, magnitude, temporal organization, oscillatory behavior, and complexity of the response.^2^ Together, they provide a multidimensional view of stimulus-induced ionic activity across different plant organs. Detailed statistical analyses are provided in the supplementary information.

### Spike Dynamics Reveal Distinct Patterns of Plant Electrical Signaling

As most existing studies characterize plant electrical responses primarily through spike number and amplitude.^2,22^ We first examined discrete spike characteristics because spike count, frequency, and amplitude provide fundamental information about the occurrence, temporal organization, and magnitude of electrical events **(Figure 2)**. Spike count revealed clear differences among stimuli and plant organs **(Figure 2a)**. Smell produced the strongest overall spike response, particularly in the root, where electrical activity was markedly elevated compared with the other stimulus conditions. This pronounced response may reflect sustained activation of root tissues during chemical stimulation.

**Figure 2:**
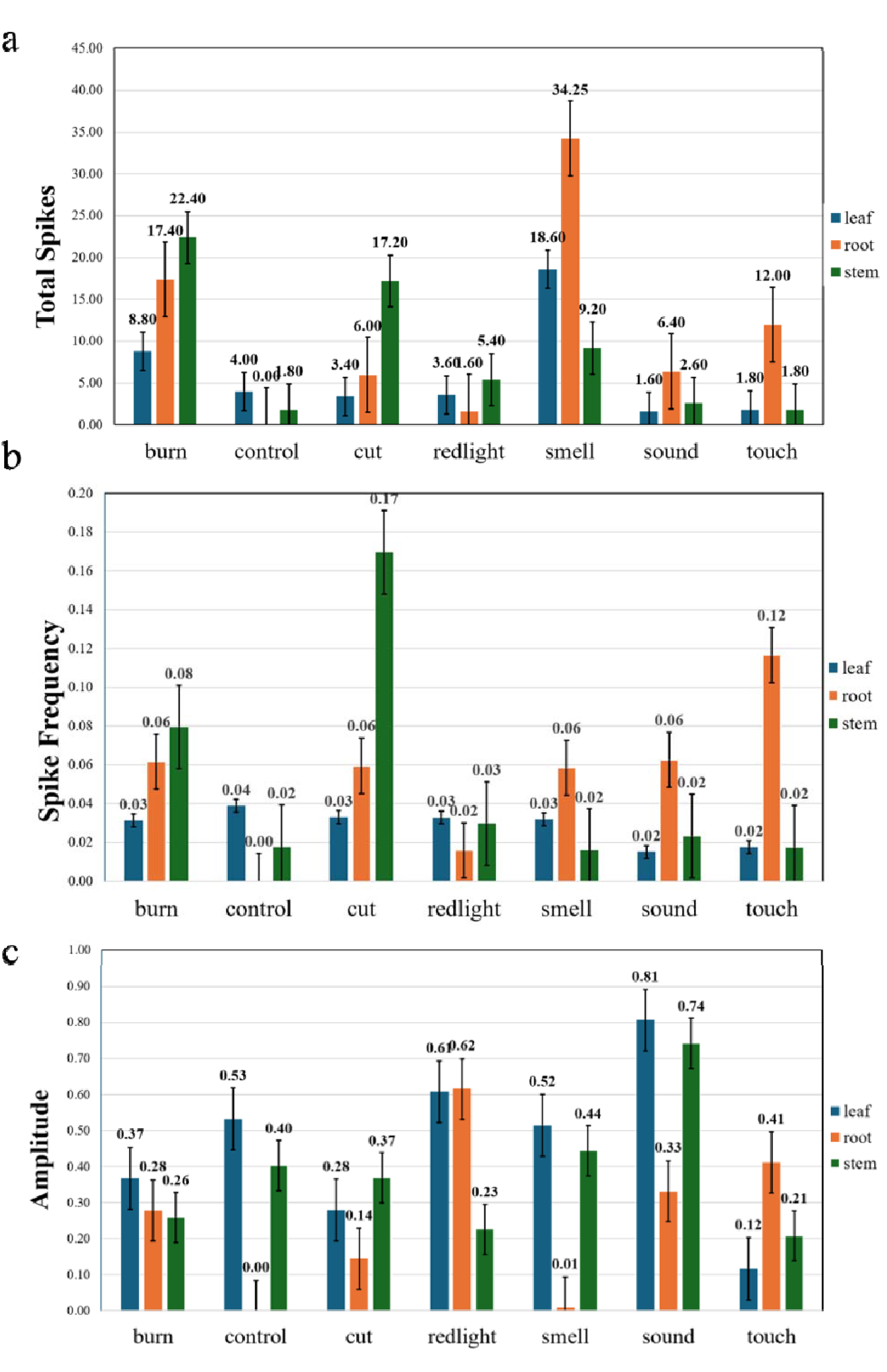
Spike-based features across stimuli and plant organs. (a) Total spike count, (b) spike frequency, and (c) mean spike amplitude for leaf, root, and stem. Strong stimuli (burn, cut, smell) show higher spike activity, while mild stimuli (touch, red light) exhibit lower responses, highlighting stimulus-dependent electrophysiological variation across organs.

Burn also produced substantial electrical activity, particularly in the stem and root, indicating broader engagement of plant tissues following thermal injury.^25^ In contrast, the leaf response remained comparatively modest. Cutting produced a predominantly stem-associated pattern, consistent with the direct mechanical disturbance experienced by the stem. Sound, touch, and red light produced comparatively limited changes in total spike production.^27^

Spike frequency revealed a pattern that was not always consistent with total spike count **(Figure 2b)**. Cutting produced the most pronounced increase in firing frequency, particularly in the stem, suggesting rapid and repeated electrical activation at or near the site of mechanical injury. Touch also produced an increase in root spike frequency despite relatively limited overall spike production. This indicates that a small number of electrical events can nevertheless occur at a relatively high rate. Burn, in contrast, produced a broader increase in frequency across the different organs, consistent with the systemic nature of the response to severe injury. Amplitude further distinguished the electrophysiological responses **(Figure 2c)**.^28,29^ Sound produced the largest individual spike amplitudes, particularly in the leaf and stem, whereas the root response was comparatively weaker. Smell showed a striking contrast between spike number and amplitude: the root generated extensive electrical activity but exhibited relatively small individual spikes. Thus, a greater number of electrical events does not necessarily indicate a greater ionic current associated with each event.^30^

Overall, no single stimulus or organ consistently dominated spike count, frequency, and amplitude. The root displayed extensive activity during smell stimulation, the stem showed rapid firing during cutting, and the leaf and stem generated large-amplitude responses to sound **(Figure 2a-c)**. These differences demonstrate that spike count, frequency, and amplitude represent distinct aspects of plant electrical signaling. Therefore, discrete-event analysis provides an important but incomplete description of stimulus-induced ionic activity, necessitating further examination of continuous waveform properties, spectral organization, and nonlinear complexity.^27–30^

### Waveform Magnitude and Structure Reflect the Plant’s Sustained Ionic Commitment

The spike-level story had answered how many cells activated, but it could not say how much energy the plant committed to sustaining that activation, or how that energy was organized in time. Continuous time-domain features, peak deflection **(Figure 3a)**, RMS amplitude **(Figure 3b)**,^31,32^ ΔAUC **(Figure 3c)**, area under the signal **(Figure 3d)**, signal variance **(Figure 3e)**, and line length **(Figure 3f)**, picked up exactly where the discrete-event picture left off, capturing the intensity, duration, and moment-to-moment structure of the ionic current itself rather than its event count **(Table S2)**.^33^

**Figure 3:**
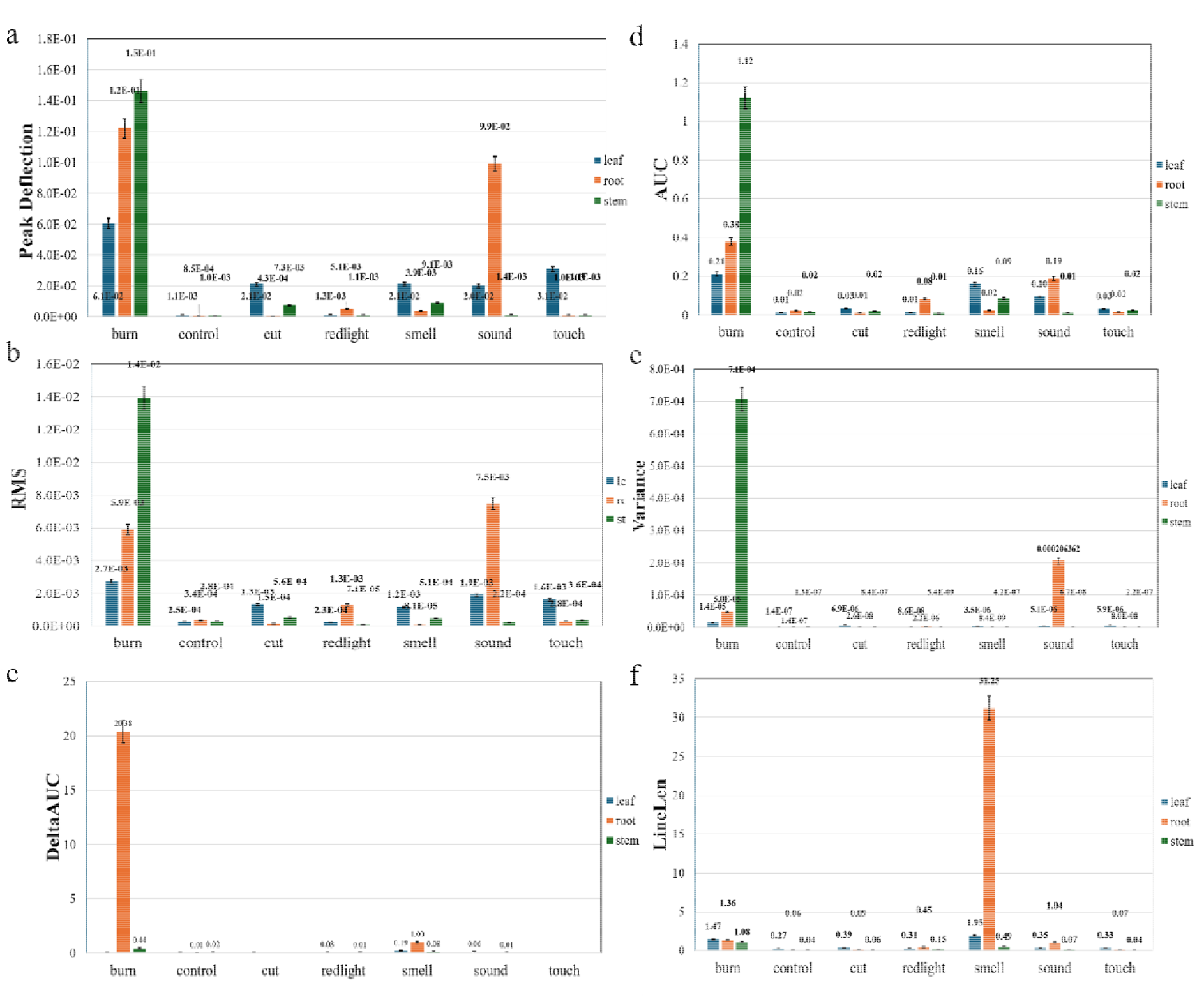
Continuous time-domain features across stimuli and plant organs (leaf, root, stem). (a) peak deflection, (b) RMS, (c) AUC, (d) ΔAUC, (e) variance, and (f) line length. Strong stimuli, particularly burn, show increased signal magnitude, energy, and variability, while mild stimuli exhibit lower and more localized responses, indicating stimulus-dependent modulation of electrical dynamics.

Burn produced the largest peak deflections and the highest RMS and AUC values of any condition, most pronounced in the stem, followed by root, with leaf showing the smallest rise of the three organs **(Figure 3a, 3b, 3d)**. This indicates that thermally induced ionic current is not only sustained but reaches its greatest magnitude in the stem, consistent with a strong hydraulic and electrical pathway running through vascular tissue rather than remaining confined to the leaf or root.^33^ Sound produced a distinct, organ-specific pattern within these same three features. Root peak deflection and RMS rose to levels approaching or exceeding those seen under burn, even though sound’s spike-level responses had been strongest in leaf and stem, showing that a stimulus can drive large, sustained root current without necessarily producing the largest discrete spike counts there. ΔAUC and variance told a sharper, more concentrated story than the other four features. ΔAUC was overwhelmingly dominated by a single condition and organ, burn in the root, where the deviation from baseline dwarfed every other organ-stimulus combination in the panel; smell produced only a modest ΔAUC rise in root and stem by comparison **(Figure 3c)**. ^34,35^ Variance showed the same concentrated pattern but in a different organ, burn drove an outsized rise in stem variance, with sound producing a smaller but still clearly elevated secondary rise in the same organ, while every other organ-stimulus pair remained close to baseline **(Figure 3e)**. These two features indicate that the most extreme, outlier-level shifts in signal trajectory and variability are concentrated in specific organs, root for ΔAUC and stem for variance, under the two most intense stimuli tested, rather than distributed evenly across the plant.

Line length broke from every pattern established so far. Rather than burn producing the broadest change, as it had for peak deflection, RMS, AUC, ΔAUC, and variance, the single largest line-length value in the entire dataset occurred in the root under smell, far exceeding root line length under any other condition, including burn **(Figure 3f)**. Burn’s line-length values, by contrast, remained comparatively modest and similar in magnitude across leaf, root, and stem. This indicates that smell drives an especially irregular, structurally complex ionic current specifically in the root, a signature not captured by any of the other five time-domain features, each of which was dominated by burn rather than smell. Taken together, the six time-domain panels show that “response magnitude” is not a single, unified axis. Burn dominates peak deflection, RMS, AUC, ΔAUC, and variance, but smell dominates line length, and it does so specifically in the root. This dissociation is itself informative, it shows that a stimulus can produce a waveform that is not especially large or energetic by most measures, yet still highly irregular in its moment-to-moment structure. Touch and red light remained the smallest responses across all six panels, consistent with locally confined ionic activity that never builds into a sustained, high-energy, or structurally complex state.^36,37^

### Spectral Composition Reflects the Rhythmic Character of Ionic Flow

Frequency-domain analysis of low-frequency band power **(Figure 4a)**, mid-frequency band power **(Figure 4b)**, and spectral entropy **(Figure 4c)** reveals which oscillatory channel of ionic activity a stimulus preferentially engages. A slow low-frequency channel associated with systemic, whole-plant ionic waves, a faster mid-frequency channel associated with secondary calcium- and reactive-oxygen-species-driven oscillations, and spectral entropy, which reflects how diverse the ionic state has become. A distal organ receiving such a signal could, in principle, distinguish the type and severity of the originating stimulus from the spectral composition of the arriving current alone, without requiring any centralized decoding step **(Table S2)**.^38^

**Figure 4:**
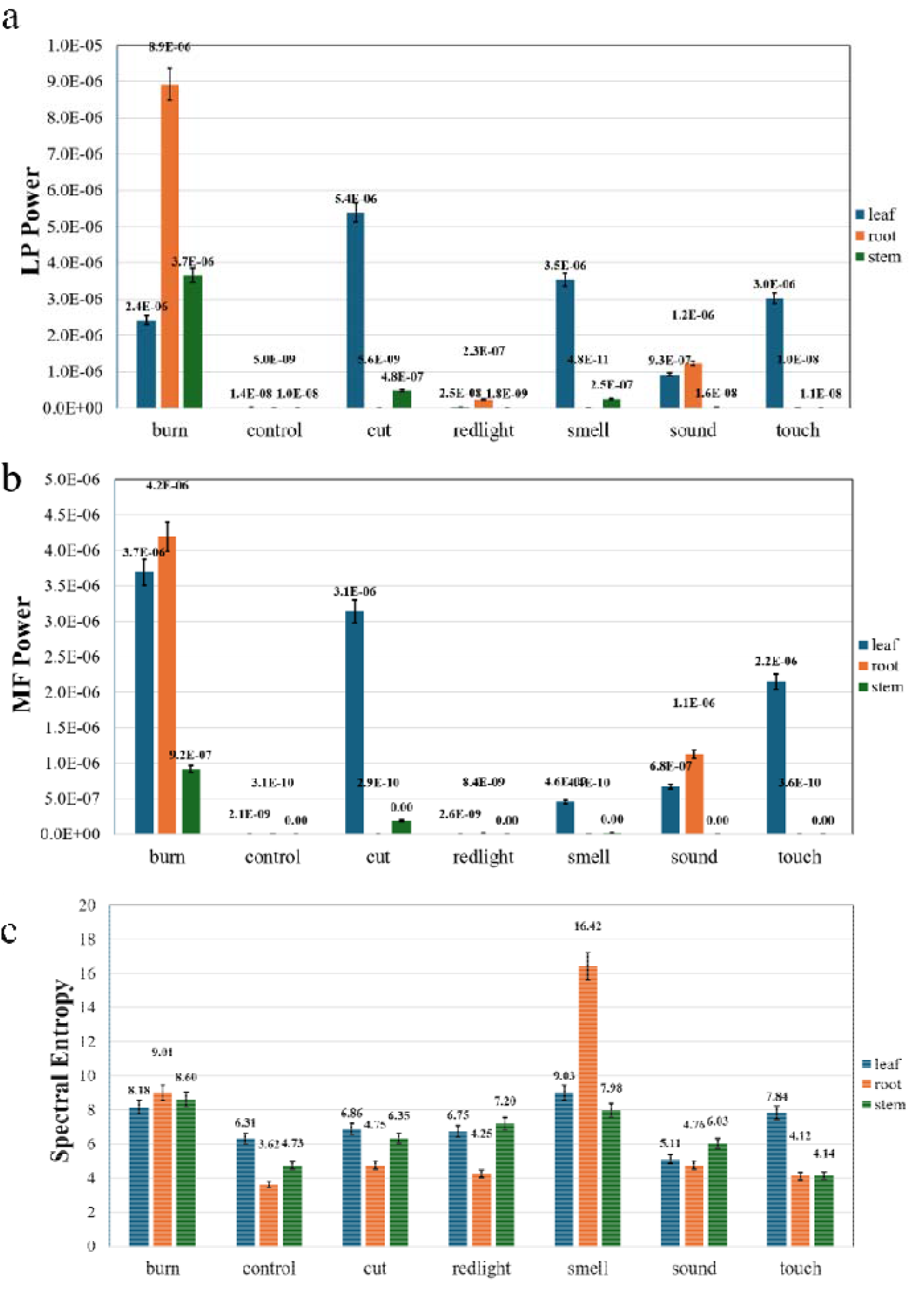
Frequency-domain features across stimuli and plant organs (leaf, stem, root). (a) low-frequency (LF) power, (b) mid-frequency (MF) power, and (c) spectral entropy. Strong and intermediate stimuli show increased spectral activity, with notable LF/MF changes indicating propagative and systemic signaling, while mild stimuli exhibit limited spectral modulation.

Burn, consistent with its dominance at every level examined so far, again produced the most comprehensive change, raising low-frequency power **(Figure 4a)**, mid-frequency power **(Figure 4b)**, and spectral entropy **(Figure 4c)** together, most extensively in the root, indicating a complete reorganization of ionic oscillatory structure under systemic thermal stress. The dominant low-frequency rise **(Figure 4a)** matches the slow, sub-hertz propagation expected of hydraulic pressure transients, while the accompanying mid-frequency rise **(Figure 4b)** reflects secondary oscillations from calcium and reactive-oxygen-species waves layered on top of that slower signal.^36^ Cutting, which had shown a trajectory-specific rather than energetic signature at the waveform level, showed the same qualitative pattern here: spectral entropy changed in the root **(Figure 4c)** without either frequency band shifting anywhere **(Figure 4a, 4b)**, reinforcing that wounding reorganizes the composition of ionic flow rather than its overall power. Sound raised low-frequency power in leaf and root but not stem **(Figure 4a)**, consistent with acoustic vibration reaching the root through the soil interface and engaging low-frequency-tuned mechanosensitive channels concentrated there.^39^ Smell, again echoing its waveform-level signature, altered spectral entropy **(Figure 4c)** and mid-frequency power **(Figure 4b)** in stem and root without raising low-frequency power anywhere **(Figure 4a)**, restructuring the qualitative character of the signal rather than adding new energy to it.^33,34^ Touch produced almost no spectral change, while red light raised both frequency bands specifically in the root **(Figure 4a, 4b)**, marking it as the exclusive site of light-induced ionic oscillation, consistent with phytochrome-mediated regulation of channel gating relayed systemically from photoreceptor cells in the leaf.^40,41^

Spectral composition tells us which oscillatory system carries a signal, but not how internally organized that ionic activity is over time, a property invisible to both energy and spectral measures alone. That gap motivated a final layer of analysis.

### Higher-Order Signal Complexity Reflects the Structural Organization of the Ionic Environment

Hjorth activity **(Figure 5a)**, mobility **(Figure 5b)**, and complexity **(Figure 5c)** capture, respectively, the variance, dominant frequency character, and structural complexity of the signal and its derivatives, chosen over entropy-, fractal-, or scaling-based measures because they remain numerically stable on short, single-trial recordings without requiring parameters difficult to justify on data of this length. Activity may reflect how energetically active the membrane ionic environment is, mobility how quickly ionic processes cycle, and complexity how many distinct ionic processes contribute at once, together characterizing the structural state of the plant’s ionic environment **(Table S2)**.^42^

**Figure 5:**
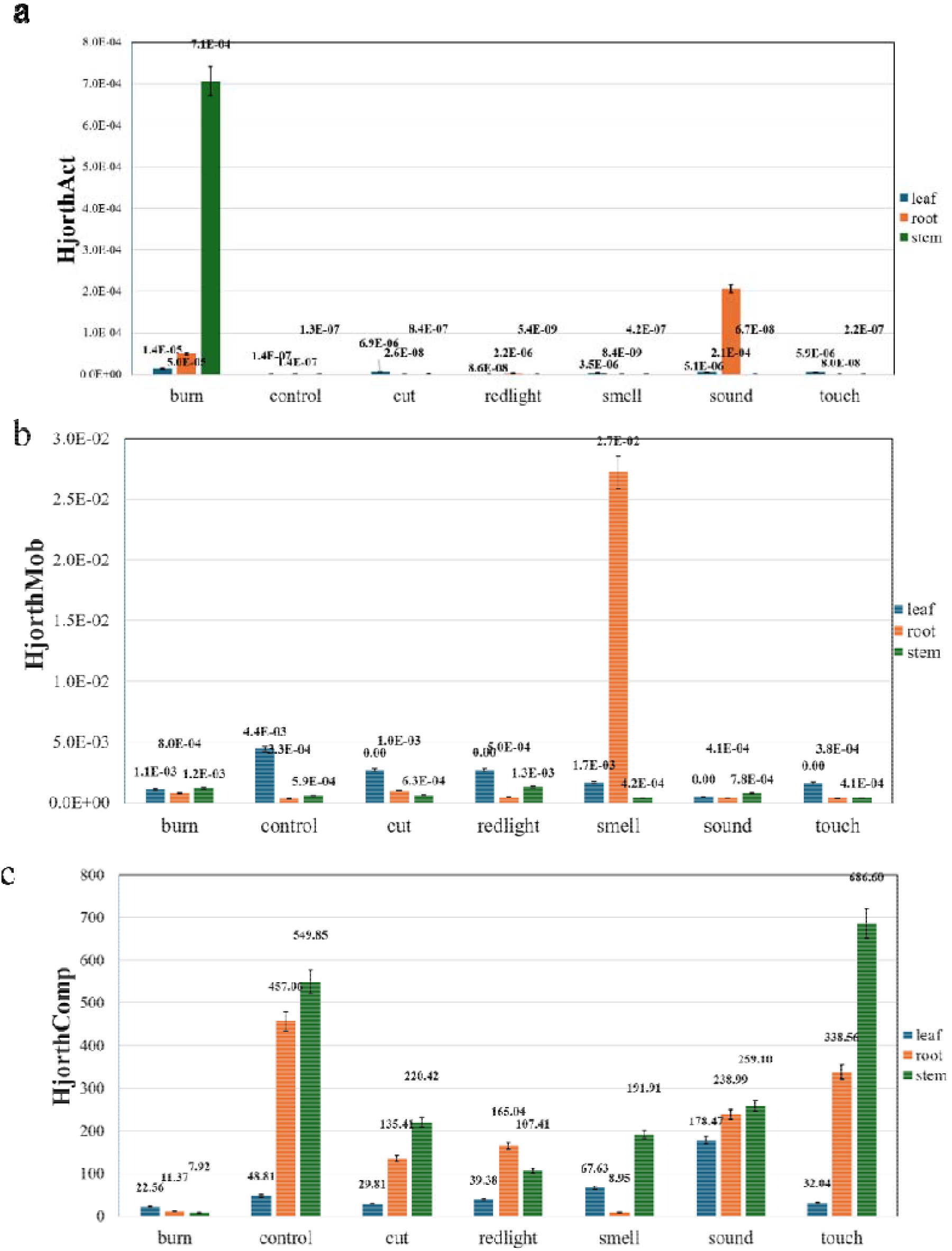
Nonlinear (Hjorth) features across stimuli and plant organs (leaf, stem, root). (a) Hjorth activity, (b) Hjorth mobility, and (c) Hjorth complexity. Strong and intermediate stimuli show increased signal dynamics and complexity, while mild stimuli exhibit relatively stable and localized responses.

Once again, burn stood apart, the root was the only organ in which all three Hjorth parameters rose together **(Figure 5a-c)**, and did so only under burn, indicating that root ionic structure is the most completely transformed of any tissue under severe stress, an outcome consistent with the simultaneous operation of several biophysical processes, elevated variance from sustained depolarization, shifted frequency character from slow-wave propagation, and added structural complexity from multiple interfering ionic sources. Cutting produced a signature here that its waveform data alone would not have predicted: Hjorth changes were confined entirely to the root **(Figure 5a-c)**, appearing despite comparatively limited time-domain change in that same tissue. This shows that mechanical wounding generates a structurally distinct ionic state at the root, arising from the convergence of vascular and mechanosensory channel activity, rather than simply a larger version of the same current, a change in the kind of signal rather than merely its size.^43,44^ Sound raised Hjorth activity broadly in leaf and root without altering complexity **(Figure 5a, 5c)**, consistent with a broadband ionic response that adds energy without introducing new structure. Smell, true to its pattern in every other domain, raised all three Hjorth parameters specifically in the root **(Figure 5a-c)** without any accompanying rise in low-frequency power, indicating chemosensory reorganization of root membrane dynamics operating below the threshold needed for detectable slow-wave accumulation. Touch and red light again produced no measurable nonlinear change anywhere **(Figure 5a-c)**, confirming across all four domains that mild stimuli leave the plant’s ionic structure largely undisturbed.^45^

### Root as a Site of Concentrated Ionic Energy Flow

By this point in the story, one pattern had recurred across every domain examined: the root, not the leaf or the stem, consistently showed the broadest and most sustained ionic engagement.^20,21^ It was the only organ in which every spike **(Figure 2a-c)**, time-domain **(Figure 3a-e)**, spectral **(Figure 4a-c)**, and nonlinear **(Figure 5a-c)** feature rose together under burn. The only organ showing coordinated nonlinear reorganization under cutting while leaf and stem showed none and the exclusive site of both light-induced spectral change and the largest spike response under smell. No single domain, taken alone, would have revealed this. It is only visible once the four analyses are read together as one continuous account of ionic activity across the same six stimuli. This convergence is consistent with the root functioning as a site of concentrated ionic energy flow, extending earlier proposals of the root apex as a structurally distinct convergence point, on the basis of its cellular organization and its role integrating gravitropic, hydrotropic, and mechanosensory input, into the electrical domain as well. Importantly, the pattern argues against a simpler explanation: that the root is merely a passive relay for currents generated in the leaf. Under cutting and smell, the root shows qualitative ionic reorganization entirely absent from aerial organs, meaning the root actively generates a distinct ionic state of its own, rather than passively receiving an amplified copy of something initiated elsewhere **(Table S3)**.

Reading the four domains together also clarifies what the recurring severity gradient represents the plant appears to scale the extent of ionic energy mobilization to the biological significance of the stimulus. Low-consequence stimuli such as touch or ambient light engage a narrow subset of ionic features in a single organ, while stimuli signaling tissue damage or systemic threat mobilize broad, coordinated ionic change spanning every domain and every organ measured, a graded, stimulus-proportional deployment of ionic energy rather than an all-or-nothing response.^29,31–42,44^

### Organism-Level Integration and Organ Specialization

Bringing the four domains together yields a coherent, organism-level account of plant electrical signaling as a spatially distributed system of ionic energy flow, rather than a set of independent local reflexes **(Figure 6, Table S4)**. In this account, environmental stimuli are the trigger, the movement of ions across membranes is the physical medium carrying the resulting signal. The vascular and apoplastic pathways conducting action, variation, and system potentials are the conduits linking organs, and the coordinated physiological adjustment of the whole plant is the downstream output **(Table 1)**.

**Figure 6:**
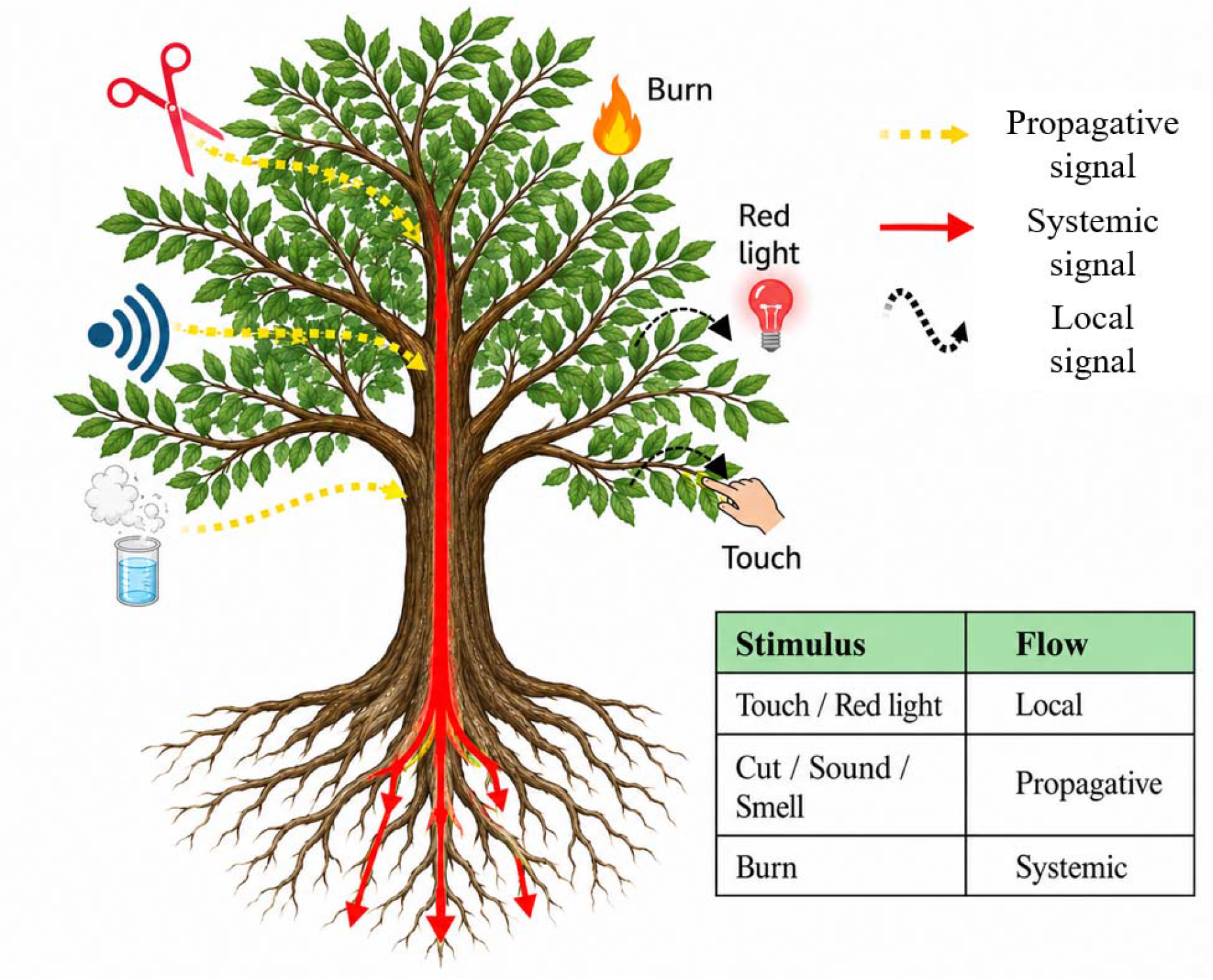
Conceptual model of stimulus-dependent hierarchy in plant electrical signaling. Mild stimuli (touch, red light) induce localized responses, intermediate stimuli (cut, sound, smell) produce propagative signaling, and strong stimuli (burn) generate systemic, long-distance integration across the plant.

Across the four domains, the six stimuli separated into three response regimes according to the extent of ionic signal propagation. Touch and red light produced predominantly local responses, with touch showing small ΔAUC and line-length changes near the stimulation site, while red light produced mainly root-specific spectral changes without broad energetic or structural reorganization. Cut and sound produced intermediate, propagative responses, engaging selected conduction routes rather than the entire plant. Cutting generated a strong localized frequency signature in the stem together with a nonlinear response in the root, consistent with transmission through vascular tissues. Sound produced a predominantly leaf-associated response with partial root involvement, suggesting propagation through mechanically responsive tissues followed by secondary transmission. Smell and burn produced systemic responses, engaging multiple organs and domains. Smell produced particularly strong root involvement, including elevated spike count, spectral entropy, and waveform irregularity, whereas burn generated the most extensive response across organs and features.

This local-propagative-systemic organization can be interpreted within the established framework of plant electrical signaling. Action potentials (APs) provide rapid electrical responses, particularly near stimulation sites, whereas variation potentials (VPs) are slower responses associated with wounding, hydraulic changes, and calcium-dependent signaling and can propagate over longer distances. System potentials (SPs) provide broader electrical coordination across tissues. These modes are not mutually exclusive and may interact with calcium and reactive-oxygen-species waves to generate stimulus-specific responses.^38,46^

At the organ level, the root frequently emerged as a site of strong signal convergence, particularly under systemic stresses, although this was stimulus-dependent.^47^ The stem primarily functioned as a transmission route, with vascular regions potentially contributing to signal propagation. The leaf predominantly acted as an environmental sensing interface, particularly for light, mechanical, and volatile stimuli, while also becoming a major response site under some conditions. Thus, the plant appears to distribute perception, transmission, and signal convergence dynamically across organs, enabling stimulus-specific responses without requiring a single centralized control site.^2^

## Conclusion

This study correlated plant bioelectrical signals with potential ionic energy flow across leaf, stem, and root of tomato (*Solanum lycopersicum*) under six stimuli, using a multi-domain framework combining spike events, waveform structure, spectral composition, and nonlinear complexity. No single organ or feature captured the full response: burn produced systemic engagement across all organs and domains, smell produced the most extreme root-specific responses, cut and sound produced propagative, organ-selective responses, and touch and red light remained largely local. This local-propagative-systemic gradient mapped onto the three classical modes of plant electrical communication, action, variation, and system potentials, recruited differentially by stimulus severity. The root showed recurring but not universal convergence, with leaf dominating several features under cut, touch, and smell. These findings support plant electrical signaling as a distributed, ion-mediated information system, with relevance for real-time stress monitoring, precision agriculture, and bio-inspired sensing design.

## Author Contributions

SC conceived and designed the experiments with inputs from CKN and LB, optimized experimental protocols, performed all plant-related experiments, data analysis, and also wrote the manuscript. CKN and LB supervised the overall project, offered continuous guidance throughout the research, and contributed to the conceptual development and editing of the manuscript.

## Data Availability Statement

The data that support the findings of this study are available in the supplementary material of this article.

## Conflict of Interest

The authors declare no conflict of interest.

## Acknowledgements

The authors acknowledge the facilities and technical assistance of the Indian Institute of Technology Mandi and the support of the IKSMHA Centre. We thank AMRC for the facility for instrument access.

## Notes

### Competing Interest Statement

The authors have declared no competing interest.

