## Supplementary Information for "Correlation of Plant Bioelectrical Signals with Potential Ionic Energy Flow under Different Stress"

### Supporting Information

#### Materials and Methods

##### Plant Material and Growth Conditions

Tomato (*Solanum lycopersicum*) seedlings of the same cultivar and age (6 weeks) were used throughout the study to minimize developmental variability. Plants were grown individually in pots containing a standardized soil mix and maintained in a controlled growth chamber at 23-25 °C, 50-60% relative humidity, under a 16 h light/8 h dark photoperiod. All plants were labeled with unique plant identifiers prior to experimentation.

To reduce physiological variability arising from hydration differences, plants were watered uniformly on the day preceding experiments, and no watering was performed within 4-6 h before recordings. Soil moisture was monitored either using a probe or by pot mass measurements and maintained within ±5% across experimental conditions.

##### Electrophysiological Recording Setup

Plant electrical activity was recorded using the Backyard Brains Plant SpikerBox system equipped with a recording electrode (clip or stake) and a ground probe. Signals were acquired using the BYB Spike Recorder software on a battery-powered laptop to minimize electrical interference from mains noise. Shielded patch cables were used throughout, and cable strain was minimized using non-metallic clips and tape.

The Plant SpikerBox system samples at approximately 10 kHz and employs an analog band-pass filter (~0.07-8.8 Hz), which is optimized for detecting plant electrical signals occurring over seconds-to-minutes timescales, including action potentials (APs), variation potentials (VPs), and slow-wave components. Signals were recorded as WAV files with synchronized event markers corresponding to stimulus onset.

**Electrode Placement and Stabilization**

The ground electrode was consistently placed in the soil at a fixed depth and location across all trials. The recording electrode was attached to the organ of interest (leaf, stem, or root), maintaining consistent inter-electrode geometry between plants and trials.

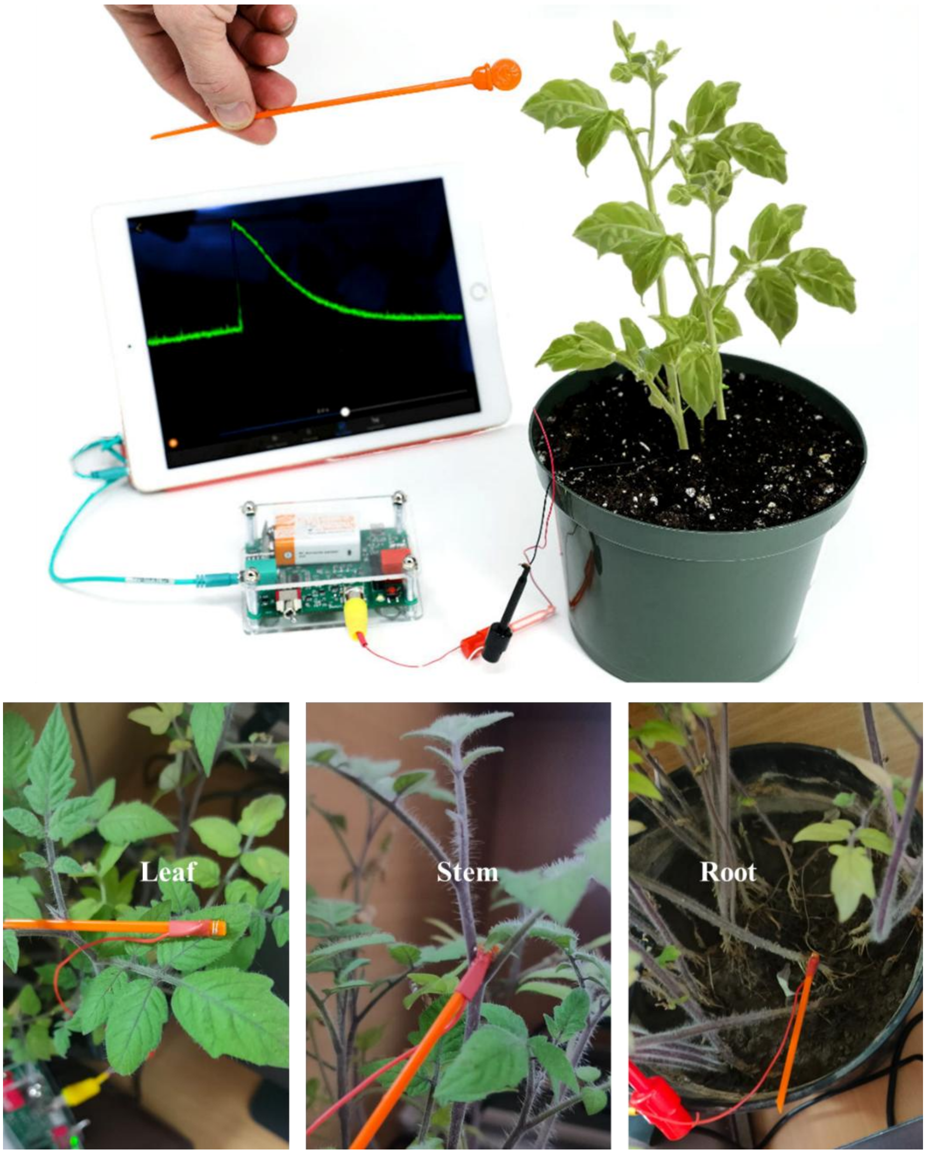


**Scheme 1:** Experimental setup for extracellular electrophysiological recording in tomato plants.

For leaf recordings, the electrode was positioned near the midrib or petiole while avoiding major veins when possible, to minimize motion artifacts. Stem recordings were performed at mid-internode regions using gentle attachment to avoid tissue compression. For root recordings, a shallow surface root was carefully exposed without damage, and electrode contact was stabilized using a damp cotton interface before re-covering to prevent desiccation. Soil-only recordings were performed with both electrodes inserted into the soil at a fixed spacing to serve as environmental controls.

After electrode placement, a stabilization period of 2-3 min was allowed to ensure baseline signal stability prior to data acquisition.

**Experimental Design and Stimuli**

| Stimulus | Stimulus | Post-stimulus | Rationale |
|---|---|---|---|
| Touch | 10-30 s | 2 min | APs within seconds; recover in minutes. |
| Cut | instant | 2 min | Rapid APs plus short slow component. |
| Burn | 2 s | 5 min | VP/slow wave lasts minutes. |
| Sound | 60 s | 2 min | Fast mechano-responses; recovery minutes. |
| Smell | 60-120 s | 10 min | Slow depolarization; VP-like. |
| Red light | 60 s | 2 min | Rapid electrical change at light step. |

Recordings were conducted from three plant organs (leaf, stem, and root). Stimuli included mechanical touch, cutting, thermal burn, sound, volatile chemical exposure (smell), and red-light illumination. For each stimulus, appropriate sham controls were implemented, including handling controls, silent speaker placement, and heat-near-leaf controls.

Mechanical touch was applied using a calibrated von Frey filament or fixed-mass probe. Cutting was performed using a sterile scalpel or micro-scissors with controlled cut length. Thermal burn stimuli were delivered using a temperature-controlled soldering iron with a blunt tip, chosen to provide reproducible heat exposure while avoiding open flame. Sound stimuli were generated using a small speaker driven by a tone generator, with sound pressure levels monitored using an SPL meter or equivalent application. Volatile chemical stimuli were delivered using acetone (or selected VOC) from a sealed vial with gentle airflow applied via tubing and syringe or pump. Red light stimulation employed a 660 nm LED source with known irradiance, with optional far-red (730 nm) illumination.

**Replication and Temporal Controls**

Experiments were conducted using a minimum of five plants per organ × stimulus combination in a between-plant design. When within-plant repetition was required, no more than three trials per day were performed on distinct sites of the same plant, with washout periods of at least 20-30 min between trials. All experiments were performed within a fixed Zeitgeber time window (ZT3-ZT6) to control for circadian influences on plant electrical signaling.

**Recording Windows and Signal Timescales**

Recording durations were selected based on the known physiological timescales of plant electrical signals. Short recording windows (2-3 min total, including baseline and post-stimulus periods) were used for stimuli such as touch, sound, and small cuts, which predominantly elicit rapid action potentials that occur within seconds and decay within minutes.

Longer recording windows (≥10 min post-stimulus) were used for burn treatments, which are known to induce variation potentials and systemic electrical waves characterized by slow depolarization and recovery phases lasting several minutes. These longer recordings were necessary to capture the full onset, peak, and decay of VP-dominated responses.

This stimulus-specific recording strategy was chosen to avoid unnecessary baseline drift while ensuring capture of biologically relevant signal components.

**Data Analysis and Feature Extraction**

Electrical recordings were analyzed using a feature-based framework designed to capture both discrete event dynamics and continuous waveform properties. Feature extraction was performed on post-stimulus segments, with baseline normalization applied where appropriate.

Event-based features included spike frequency, mean absolute spike amplitude, and the coefficient of variation of inter-spike intervals (ISI CV). These metrics were selected to characterize event rate, strength, and temporal organization while avoiding over-reliance on raw spike counts.

Continuous time-domain features included peak absolute deflection, root-mean-square (RMS) amplitude, area under the rectified signal (AUC), signal variance, and line length, providing robust measures of response magnitude, duration, and overall electrical activity. Frequency-domain features comprised low-frequency (0.1-1 Hz) and mid-frequency (1-5 Hz) band power, along with spectral entropy, to quantify variation potential-dominated signaling and spectral organization. Nonlinear signal structure was assessed using Hjorth activity, mobility, and complexity parameters.

To account for inter-plant variability and baseline drift, stimulus-evoked changes were quantified using baseline-normalized ΔAUC (post-stimulus minus pre-stimulus), providing a within-trial measure of stimulus-specific electrical response.

**Statistical Analysis**

Extracted features were statistically compared across control and stimulus conditions, and across organs, using appropriate parametric or non-parametric tests depending on data distribution. Analysis of variance (ANOVA) was used for multi-group comparisons, followed by pairwise comparisons where applicable. All statistical analyses were performed on per-trial feature values to enable robust comparison across experimental conditions. All statistical analyses were performed on the original feature values. Because a large number of feature, organ, and stimulus combinations were tested, the reported p-values are not corrected for multiple comparisons and are provided for descriptive purposes; multiple-comparison correction and effect-size reporting are addressed in the Limitations section.

**Safety Considerations**

All experiments involving heat, and volatile chemicals were conducted using appropriate safety precautions, including heat-resistant surfaces, fume extraction, gloves, and eye protection.

#### Supplementary Tables

All tables are provided here as supplementary material. They are referenced in the main text as Table S1 to Table S4.

**Table S1.** Correspondence between general information-processing functions and the electrophysiological processes observed in plant electrical signaling in this study. Column 3 reports what was directly measured; column 4 reports the established biophysical basis, drawn from prior literature rather than independently verified here (no calcium imaging, ion-selective electrodes, or channel blockers were used in this study).

| Functional Process | General Description | Directly Observed in This Study | Established Ionic Basis (from prior literature) |
|---|---|---|---|
| Detection | A physical change occurs at the site of stimulation | Statistically distinct electrical responses recorded at the stimulated organ for all six stimuli | Mechanosensitive, thermosensitive, and ligand-gated ion channels open in response to the corresponding stimulus class |
| Stimulus Differentiation | Different stimuli produce different physical signatures | Distinct spike, time-domain, frequency, and nonlinear profiles per stimulus condition | Different stimuli are known to preferentially engage different channel populations (e.g., mechanosensitive vs. thermosensitive channels) |
| Signal Propagation | A local signal is transmitted to distant tissue | Organ-specific signals recorded in leaf, stem, and root confirm inter-organ signal transmission | Action potentials propagate via sequential voltage-gated channel activation; variation potentials propagate via hydraulic pressure waves coupled to channel activation |
| Convergence | Signals from multiple sources arrive together at one site | Root shows the broadest multi-feature response across all stimuli in this dataset | Not independently established in this study; consistent with prior anatomical proposals of the root apex as an integration zone |
| Downstream Coupling | An electrical signal is associated with a subsequent physiological process | Not measured directly in this study | Established in other systems: plant electrical signals are known to be temporally coupled with $Ca^{2+}$ waves, ROS bursts, and hormonal signaling |
| Physiological Outcome | The organism's physiological state changes following signal processing | Not measured directly in this study | Prior work links electrical signaling to defense gene expression, stomatal closure, and systemic wound responses |

**Table S2.** Time-domain, frequency-domain, and nonlinear features: direct measurement and established interpretation.

| Feature | Direct Measurement | Established Interpretation |
|---|---|---|
| Spike Count* | Discrete event count | Each event is consistent with a depolarization event at or near the recording site, most likely reflecting local ion channel activation |
| Spike Frequency* | Event rate | May reflect the rate at which depolarization events are triggered or relayed; not a direct measure of single-channel gating kinetics |
| Spike Amplitude* | Peak deflection per event | May reflect the magnitude of net ionic current at the electrode; extracellular amplitude is influenced by electrode placement and tissue geometry as well as underlying current magnitude |
| Peak Deflection | Maximum voltage displacement from baseline | Reflects the largest net signal recorded in the window; not attributable to a specific channel type without pharmacological or imaging validation |
| RMS | Root-mean-square of the signal amplitude | A standard measure of overall signal energy; commonly used as a proxy for the intensity of the electrical response in plant electrophysiology |
| AUC | Area under the rectified signal | A standard measure of cumulative signal magnitude over time |
| ΔAUC | Change in AUC relative to pre-stimulus baseline | Quantifies stimulus-evoked deviation independent of absolute signal magnitude |
| Variance | Statistical variance of the signal | A standard descriptor of signal variability; does not by itself indicate the number of contributing sources |
| Line Length | Cumulative absolute difference between successive samples | A standard measure of waveform irregularity, used elsewhere in EEG and plant electrophysiology as an index of signal complexity |
| LF Power** | Power in the 0.1–1 Hz band | This frequency range is consistent with the known timescale of variation-potential propagation reported in the plant electrophysiology literature |
| MF Power** | Power in the 1–5 Hz band | This frequency range overlaps with oscillation frequencies reported for calcium- and ROS-associated signaling in other plant studies; not independently confirmed here |
| Spectral Entropy** | Shannon entropy of the power spectrum | A standard measure of spectral complexity/diversity; higher entropy indicates power spread across more frequency components |
| Hjorth Activity*** | Variance of the raw signal | Standard time-domain complexity parameter; indexes overall signal power |
| Hjorth Mobility*** | Ratio of the standard deviation of the first derivative to that of the signal | Standard parameter; approximates the mean frequency of the signal |
| Hjorth Complexity*** | Ratio of the mobility of the first derivative to the mobility of the signal | Standard parameter; approximates deviation from a pure sine wave, i.e., waveform irregularity |

** Spike-based domain. ** Frequency domain. *** Nonlinear domain.*

**Table S4.** Stimulus-dependent pattern of electrophysiological engagement across domains, as directly observed in this study.

| Stimulus | Multi-Domain Pattern Observed | Note |
|---|---|---|
| Control | No significant change in any feature or organ | Baseline recording condition |
| Red Light | Change in ΔAUC and line length in root only; LF and MF power change in root | Response confined to a single organ and a subset of features |
| Touch | Change in ΔAUC in leaf and stem; line length change in root | Response limited to trajectory-sensitive measures; no energetic or spectral change |
| Sound | Multi-feature change in leaf; LF power change in root; Hjorth activity change in leaf and root | Response engages both aerial and root tissue across several domains |
| Smell | Highest spike count of any organ-stimulus pair in root; AUC and line length change in multiple organs; Hjorth reorganization in root | Broadest single-organ engagement recorded in the spike domain |
| Cut | Organ-selective changes concentrated in stem (frequency) and root (nonlinear structure) | Root shows nonlinear change despite limited time-domain change in the same tissue |
| Burn | Significant change in nearly every feature, in every organ, across all four domains | Most extensive response recorded in this study |

**Table S5.** Structural and physiological elements of the plant electrical signaling system and their established role, as supported by this study's data or by prior literature.

| Structural or Physiological Element | Role | Basis |
|---|---|---|
| Environmental stimulus | Initiates the electrical response | Directly manipulated in this study (six stimulus types) |
| Extracellular bioelectrical activity | The measured signal | Directly recorded in this study |
| Vascular and apoplastic routes (phloem, xylem, apoplast) | Established anatomical pathways for AP, VP, and SP propagation | Established in prior plant electrophysiology literature; not independently traced in this study |
| Root apex | Organ showing the broadest and most consistent multi-domain response in this dataset | Directly observed in this study; interpretation as a "coordination center" follows prior anatomical/functional proposals, not new evidence presented here |
| Whole-plant electrical network | The organism-level system inferred by combining organ-resolved recordings | Inferred from this study's multi-organ dataset |
| Downstream physiological response (defense, hormonal, hydraulic) | Proposed physiological consequence of electrical signaling | Not measured in this study; based on established literature |

#### Statistical Analysis of Time domain, Frequency domain and non-linear

##### Control vs cut (leaf)

*Independent Samples T-Test*

| | U | df | p |
|---|---|---|---|
| DeltaAUC | 18.00 | | .31 |
| PeakDefl | 10.00 | | .69 |
| RMS | 8.00 | | .42 |
| AUC | 8.00 | | .42 |
| Variance | 8.00 | | .42 |
| LineLen | 0.00 | | < .01 |
| LFpower | 12.00 | | 1.00 |
| MFpower | 7.00 | | .31 |
| SpecEntropy | 8.00 | | .42 |
| HjorthAct | 8.00 | | .42 |
| HjorthMob | 16.00 | | .55 |
| HjorthComp | 13.00 | | 1.00 |

*Note.* Mann-Whitney U test.

##### Control vs cut (stem)

*Independent Samples T-Test*

| | U | df | p |
|---|---|---|---|
| DeltaAUC | 25.00 | | < .01 |
| PeakDefl | 6.00 | | .22 |
| RMS | 12.00 | | 1.00 |
| AUC | 15.00 | | .69 |
| Variance | 12.00 | | 1.00 |
| LineLen | 4.00 | | .10 |
| LFpower | 7.00 | | .31 |
| MFpower | 4.00 | | .10 |
| SpecEntropy | 4.00 | | .10 |
| HjorthAct | 12.00 | | 1.00 |
| HjorthMob | 8.00 | | .42 |
| HjorthComp | 18.00 | | .31 |

*Note.* Mann-Whitney U test.

**Control vs cut (root)**

*Independent Samples T-Test*

| | U | df | p |
|---|---|---|---|
| DeltaAUC | 25.00 | | < .01 |
| PeakDefl | 23.00 | | .03 |
| RMS | 23.00 | | .03 |
| AUC | 23.00 | | .03 |
| Variance | 23.00 | | .03 |
| LineLen | 0.00 | | < .01 |
| LFpower | 14.00 | | .84 |
| MFpower | 14.00 | | .84 |
| SpecEntropy | 0.00 | | < .01 |
| HjorthAct | 23.00 | | .03 |
| HjorthMob | 1.00 | | .02 |
| HjorthComp | 25.00 | | < .01 |

*Note.* Mann-Whitney U test.

**Control vs burn (leaf)**

*Independent Samples T-Test*

| | U | df | p |
|---|---|---|---|
| DeltaAUC | 10.00 | | .69 |
| PeakDefl | 0.00 | | < .01 |
| RMS | 2.00 | | .03 |
| AUC | 0.00 | | < .01 |
| Variance | 2.00 | | .03 |
| LineLen | 0.00 | | < .01 |
| LFpower | 0.00 | | < .01 |
| MFpower | 1.00 | | .02 |
| SpecEntropy | 7.00 | | .31 |
| HjorthAct | 2.00 | | .03 |
| HjorthMob | 20.00 | | .15 |
| HjorthComp | 15.00 | | .69 |

*Note.* Mann-Whitney U test.

**Control vs burn (stem)**

*Independent Samples T-Test*

| | U | df | p |
|---|---|---|---|
| DeltaAUC | 6.00 | | .22 |
| PeakDefl | 0.00 | | < .01 |
| RMS | 0.00 | | < .01 |
| AUC | 0.00 | | < .01 |
| Variance | 0.00 | | < .01 |
| LineLen | 0.00 | | < .01 |
| LFpower | 0.00 | | < .01 |
| MFpower | 0.00 | | < .01 |
| SpecEntropy | 2.00 | | .03 |
| HjorthAct | 0.00 | | < .01 |
| HjorthMob | 8.00 | | .42 |
| HjorthComp | 25.00 | | < .01 |

*Note.* Mann-Whitney U test.

**Control vs burn (root)**

*Independent Samples T-Test*

| | U | df | p |
|---|---|---|---|
| DeltaAUC | 0.00 | | < .01 |
| PeakDefl | 0.00 | | < .01 |
| RMS | 0.00 | | < .01 |
| AUC | 0.00 | | < .01 |
| Variance | 0.00 | | < .01 |
| LineLen | 0.00 | | < .01 |
| LFpower | 0.00 | | < .01 |
| MFpower | 0.00 | | < .01 |
| SpecEntropy | 0.00 | | < .01 |
| HjorthAct | 0.00 | | < .01 |
| HjorthMob | 1.00 | | .02 |
| HjorthComp | 25.00 | | < .01 |

*Note.* Mann-Whitney U test.

**Control vs touch (leaf)**

*Independent Samples T-Test*

| | U | df | p |
|---|---|---|---|
| DeltaAUC | 25.00 | | < .01 |
| PeakDefl | 4.00 | | .10 |
| RMS | 5.00 | | .15 |
| AUC | 8.00 | | .42 |
| Variance | 5.00 | | .15 |
| LineLen | 10.00 | | .69 |
| LFpower | 3.00 | | .06 |
| MFpower | 2.00 | | .03 |
| SpecEntropy | 5.00 | | .15 |
| HjorthAct | 5.00 | | .15 |
| HjorthMob | 16.00 | | .55 |
| HjorthComp | 17.00 | | .42 |

*Note.* Mann-Whitney U test.

**Control vs touch (stem)**

*Independent Samples T-Test*

| | U | df | p |
|---|---|---|---|
| DeltaAUC | 25.00 | | < .01 |
| PeakDefl | 11.00 | | .84 |
| RMS | 9.00 | | .55 |
| AUC | 10.00 | | .69 |
| Variance | 9.00 | | .55 |
| LineLen | 4.00 | | .10 |
| LFpower | 12.00 | | 1.00 |
| MFpower | 16.00 | | .55 |
| SpecEntropy | 17.00 | | .42 |
| HjorthAct | 9.00 | | .55 |
| HjorthMob | 16.00 | | .55 |
| HjorthComp | 9.00 | | .55 |

*Note.* Mann-Whitney U test.

**Control vs touch (root)**

*Independent Samples T-Test*

| | U | df | p |
|---|---|---|---|
| DeltaAUC | 30.00 | | < .01 |
| PeakDefl | 11.00 | | .54 |
| RMS | 19.00 | | .54 |
| AUC | 21.00 | | .33 |
| Variance | 19.00 | | .54 |
| LineLen | 0.00 | | < .01 |
| LFpower | 10.00 | | .43 |
| MFpower | 7.00 | | .18 |
| SpecEntropy | 2.00 | | .02 |
| HjorthAct | 19.00 | | .54 |
| HjorthMob | 9.00 | | .33 |
| HjorthComp | 23.00 | | .18 |

*Note.* Mann-Whitney U test.

**Control vs sound (leaf)**

*Independent Samples T-Test*

| | U | df | p |
|---|---|---|---|
| DeltaAUC | 4.00 | | .10 |
| PeakDefl | 2.00 | | .03 |
| RMS | 1.00 | | .02 |
| AUC | 1.00 | | .02 |
| Variance | 1.00 | | .02 |
| LineLen | 0.00 | | < .01 |
| LFpower | 0.00 | | < .01 |
| MFpower | 4.00 | | .10 |
| SpecEntropy | 19.00 | | .22 |
| HjorthAct | 1.00 | | .02 |
| HjorthMob | 23.00 | | .03 |
| HjorthComp | 4.00 | | .10 |

*Note.* Mann-Whitney U test.

**Control vs sound (stem)**

*Independent Samples T-Test*

| | U | df | p |
|---|---|---|---|
| DeltaAUC | 18.00 | | .31 |
| PeakDefl | 9.00 | | .55 |
| RMS | 13.00 | | 1.00 |
| AUC | 15.00 | | .69 |
| Variance | 13.00 | | 1.00 |
| LineLen | 0.00 | | < .01 |
| LFpower | 9.00 | | .55 |
| MFpower | 8.00 | | .42 |
| SpecEntropy | 6.00 | | .22 |
| HjorthAct | 13.00 | | 1.00 |
| HjorthMob | 9.00 | | .55 |
| HjorthComp | 18.00 | | .31 |

*Note.* Mann-Whitney U test.

**Control vs sound (root)**

*Independent Samples T-Test*

| | U | df | p |
|---|---|---|---|
| DeltaAUC | 20.00 | | .15 |
| PeakDefl | 2.00 | | .03 |
| RMS | 2.00 | | .03 |
| AUC | 3.00 | | .06 |
| Variance | 2.00 | | .03 |
| LineLen | 0.00 | | < .01 |
| LFpower | 0.00 | | < .01 |
| MFpower | 4.00 | | .10 |
| SpecEntropy | 4.00 | | .10 |
| HjorthAct | 2.00 | | .03 |
| HjorthMob | 12.00 | | 1.00 |
| HjorthComp | 20.00 | | .15 |

*Note.* Mann-Whitney U test.

**Control vs smell (leaf)**

*Independent Samples T-Test*

| | U | df | p |
|---|---|---|---|
| DeltaAUC | 0.00 | | < .01 |
| PeakDefl | 4.00 | | .10 |
| RMS | 7.00 | | .31 |
| AUC | 2.00 | | .03 |
| Variance | 7.00 | | .31 |
| LineLen | 0.00 | | < .01 |
| LFpower | 8.00 | | .42 |
| MFpower | 4.00 | | .10 |
| SpecEntropy | 4.00 | | .10 |
| HjorthAct | 7.00 | | .31 |
| HjorthMob | 17.00 | | .42 |
| HjorthComp | 10.00 | | .69 |

*Note.* Mann-Whitney U test.

**Control vs smell (stem)**

*Independent Samples T-Test*

| | U | df | p |
|---|---|---|---|
| DeltaAUC | 2.00 | | .03 |
| PeakDefl | 2.00 | | .03 |
| RMS | 6.00 | | .22 |
| AUC | 0.00 | | < .01 |
| Variance | 6.00 | | .22 |
| LineLen | 0.00 | | < .01 |
| LFpower | 4.00 | | .10 |
| MFpower | 0.00 | | < .01 |
| SpecEntropy | 0.00 | | < .01 |
| HjorthAct | 6.00 | | .22 |
| HjorthMob | 10.00 | | .69 |
| HjorthComp | 18.00 | | .31 |

*Note.* Mann-Whitney U test.

**Control vs smell (root)**

*Independent Samples T-Test*

| | U | df | p |
|---|---|---|---|
| DeltaAUC | 0.00 | | .02 |
| PeakDefl | 6.00 | | .41 |
| RMS | 20.00 | | .02 |
| AUC | 8.00 | | .73 |
| Variance | 20.00 | | .02 |
| LineLen | 0.00 | | .02 |
| LFpower | 20.00 | | .02 |
| MFpower | 15.00 | | .29 |
| SpecEntropy | 0.00 | | .02 |
| HjorthAct | 20.00 | | .02 |
| HjorthMob | 0.00 | | .02 |
| HjorthComp | 20.00 | | .02 |

*Note.* Mann-Whitney U test.

**Control vs redlight (leaf)**

*Independent Samples T-Test*

| | U | df | p |
|---|---|---|---|
| DeltaAUC | 10.00 | | .69 |
| PeakDefl | 12.00 | | 1.00 |
| RMS | 11.00 | | .84 |
| AUC | 9.00 | | .55 |
| Variance | 11.00 | | .84 |
| LineLen | 7.00 | | .31 |
| LFpower | 9.00 | | .55 |
| MFpower | 5.00 | | .15 |
| SpecEntropy | 10.00 | | .69 |
| HjorthAct | 11.00 | | .84 |
| HjorthMob | 14.00 | | .84 |
| HjorthComp | 12.00 | | 1.00 |

*Note.* Mann-Whitney U test.

**Control vs redlight (stem)**

*Independent Samples T-Test*

| | U | df | p |
|---|---|---|---|
| DeltaAUC | 13.00 | | 1.00 |
| PeakDefl | 10.00 | | .69 |
| RMS | 21.00 | | .10 |
| AUC | 17.00 | | .42 |
| Variance | 21.00 | | .10 |
| LineLen | 8.00 | | .42 |
| LFpower | 17.00 | | .42 |
| MFpower | 12.00 | | 1.00 |
| SpecEntropy | 3.00 | | .06 |
| HjorthAct | 21.00 | | .10 |
| HjorthMob | 4.00 | | .10 |
| HjorthComp | 21.00 | | .10 |

*Note.* Mann-Whitney U test.

**Control vs redlight (root)**

*Independent Samples T-Test*

| | U | df | p |
|---|---|---|---|
| DeltaAUC | 25.00 | | < .01 |
| PeakDefl | 3.00 | | .06 |
| RMS | 4.00 | | .10 |
| AUC | 5.00 | | .15 |
| Variance | 4.00 | | .10 |
| LineLen | 0.00 | | < .01 |
| LFpower | 1.00 | | .02 |
| MFpower | 1.00 | | .02 |
| SpecEntropy | 4.00 | | .10 |
| HjorthAct | 4.00 | | .10 |
| HjorthMob | 7.00 | | .31 |
| HjorthComp | 24.00 | | .02 |

*Note.* Mann-Whitney U test.
